# Context-Dependent Modulations in Acoustic Features of Zebra Finch Distance Calls: Insights from a Novel Goal-Directed Vocalization Paradigm

**DOI:** 10.1101/2024.09.24.614738

**Authors:** Zohreh Safarcharati, Amirreza Bahramani, Pouya Mokari Amjad, Mahsa Ravanbakhsh, Mohammad Reza Raoufy, Mahdi Khademian

**Affiliations:** School of Cognitive Sciences, Institute for Research in Fundamental Sciences (IPM), Tehran 1956836484, Iran; Department of Cognitive Neuroscience, Faculty of Interdisciplinary Sciences and Technologies, Tarbiat Modares University, Tehran, Iran; Department of Electrical Engineering, Sharif University of Technology, Tehran 1458889694, Iran; Sharif Brain Center, Sharif University of Technology, Tehran 1458889694, Iran; Department of Physiology, Faculty of Medical Sciences, Tarbiat Modares University, Tehran, Iran; Institute for Brain Sciences and Cognition, Faculty of Medical Sciences, Tarbiat Modares University, Tehran, Iran

**Keywords:** Vocal Communication, Zebra Finch, Distance Call, Operant Conditioning, Vocalization Paradigm, Acoustic Flexibility

## Abstract

Songbirds are renowned for their complex vocal communication abilities; among them, zebra finches (*Taeniopygia guttata*) are a key species for studying vocal learning and communication. Zebra finches use various calls with different meanings, including the distance call, which is used for long-distance contact. Whether these calls are static with fixed meanings or flexible remains an open question. In this study we aimed to answer this question by designing a novel behavioral paradigm, in which we trained food-restricted zebra finches to use distance calls for food request. Nine out of ten birds learned this association and used their distance calls to obtain food when they were hungry. We then introduced a visually-separated audience and compared the distance calls used for food requests with those used for communication between birds. Results revealed significant acoustic differences in power, pitch, and other spectral characteristics between the distance calls uttered in these two contexts. Our findings suggest that zebra finches can use their distance call for different goals and also acoustically modulate it based on the context. Therefore, it demonstrates a level of vocal control thought to be exclusive to songs. This study enhances our understanding of vocal flexibility and its role in vocal communication.

## Introduction

Vocal communication plays a fundamental role in social interactions, being crucial for survival and other aspects of life in many animal species ^1^. Songbirds, in particular, are well known for their ability to learn and produce complex songs, a trait that has made them key models for investigating the behavioral and neural mechanisms of vocal learning and communication ^2^. Among vast species of songbirds, zebra finches (*Taeniopygia guttata*) have been the most popular animal model for many reasons especially the simplicity of their vocalizations ^3^. Although much research has focused on their singing behavior, the significance of their calls—shorter and less complex vocalizations—has been comparatively overlooked ^4^. Calls serve various functions in the social and environmental contexts of songbirds, including mating interactions, alerting conspecifics to danger, coordinating activities such as nest building and so forth ^3^.

Among the various types of calls, distance calls are of particular interest. These calls are mainly produced when birds are visually separated and trying to maintain contact, but they are also used during take-off, greetings, between bouts of singing and etc. ^3^. Another important aspect is that the male distance call is the only learned call among zebra finch call repertoire ^5^. This characteristic leads to structural variations in the distance calls across different zebra finches^6–8^.

Recent studies have begun to unveil the complexity and plasticity of songbird calls, challenging the earlier notion that these vocalizations are simple, invariant and involuntary responses. For instance some internal states such as stress ^9^ and access to water ^10^ can influence the acoustic features and dynamics of calls. Furthermore zebra finches can use the acoustic features of calls to recognize their mates ^11,12^, their parents ^13^ and some other conspecific individuals ^14^. These findings suggest that calls, like songs ^15^, possess a degree of flexibility and may convey more information than previously assumed. This raises an important question: Are calls fixed in meaning, or can they be adapted for different functions depending on the context?

To investigate this, traditional behavioral paradigms that rely on operant conditionings, such as key-pressing assays ^16–18^ or operant discrimination tasks ^19–23^ are insufficient. While these methods are valuable for many behavioral studies, they do not require the birds to use their vocalizations, which is essential for studying the flexibility and contextual use of calls. Recently, novel paradigms have been developed to approach this problem in which the bird was instructed to initiate vocalization ^24,25^, but to the best of our knowledge none have been designed specifically for zebra finches. Thus, a new paradigm is needed to address these questions.

In this study, we present the design of a novel behavioral paradigm that trains food-restricted zebra finches to use their distance calls to obtain food. For our paradigm to be effective, we developed a robust and stable closed-loop vocalization detection and classification system. This system employs a vocal activity detection (VAD) algorithm to reliably identify vocalizations, which are then classified in real-time.

We interestingly observed that the birds had the ability to learn using their distance calls for food request. After that we investigated whether these calls differed from those used for long-distance contact. Importantly, our findings revealed significant differences in the acoustic features of distance calls associated with food acquisition compared to those used for responding to conspecific distance calls. Specifically, distance calls associated with feeding exhibited distinct changes in power, pitch, and some other spectral characteristics. These findings suggest that zebra finches can modulate their calls in a goal-directed manner, providing further evidence of call plasticity and contextual flexibility. This challenges the traditional view that calls are fixed in meaning and suggests that they can be adapted for different goals.

## Results

### Closed-loop VAD system for goal-directed vocalization paradigm

The goal of this paradigm was to train subjects to use specific vocalizations to obtain food. This required real-time analysis of the birds’ vocalizations, identifying the appropriate ones, and using a closed-loop VAD system to trigger the feeder when the correct vocalization was detected (Fig. 1).

**Figure 1.**
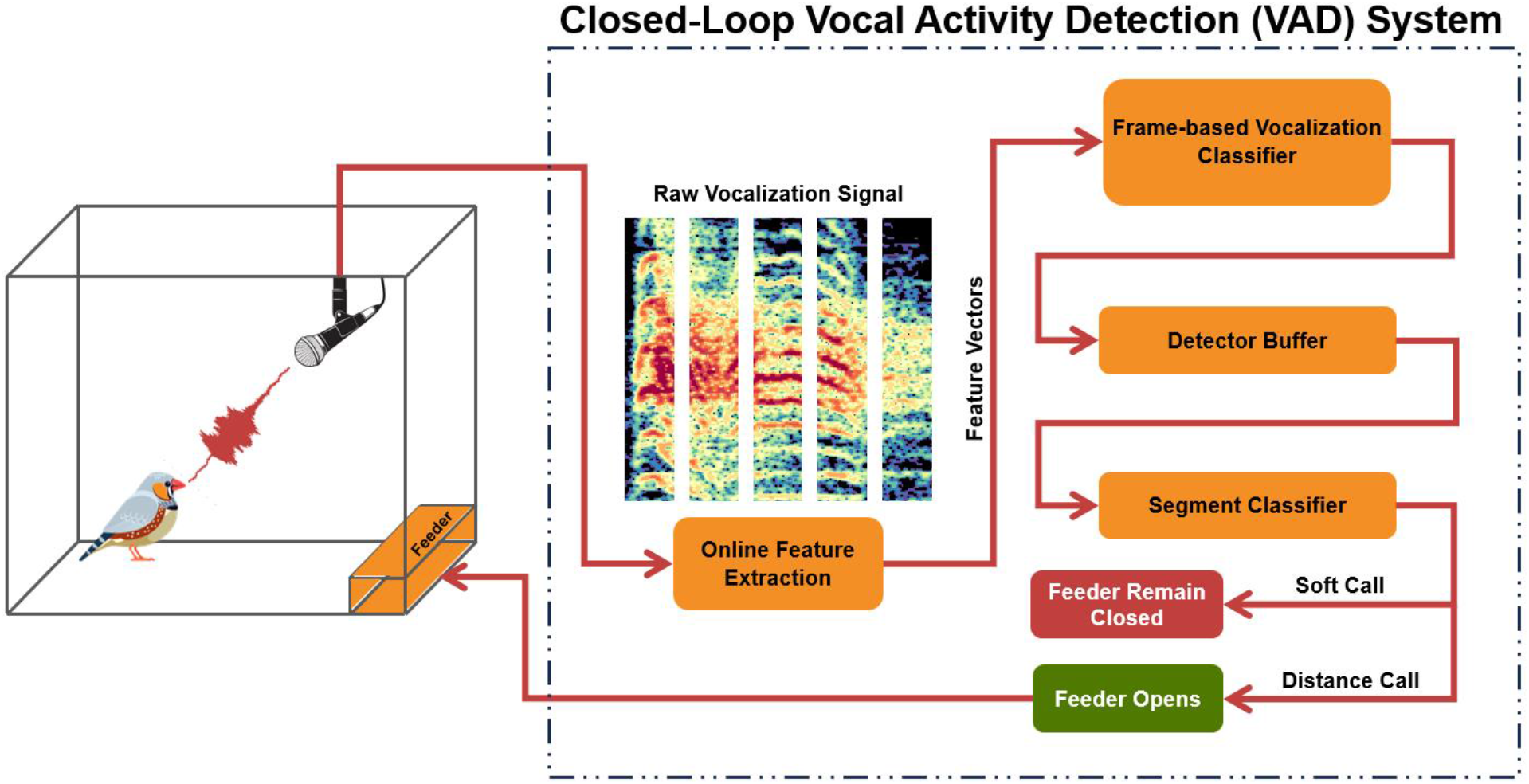
Schematic of the experimental setup and closed-loop Vocal Activity Detection (VAD) system architecture. Initially, the microphone records the bird vocalizations and transmits the signal to the control computer. The VAD system then segments the vocalization into overlapping windows. Acoustic features are extracted from each part and the whole vocalization is classified as either a “distance call” or “other calls”. If the vocalization was identified as a distance call, the feeder automatically opens and remained open for five seconds, allowing the bird to access food.

To achieve this, we implemented a closed-loop VAD system. Although various methods have been used previously for this purpose ^26–29^, we aimed to develop a system that was more robust to environmental noise. This VAD system utilizes a unidirectional Long Short-Term Memory (LSTM) Recurrent Neural Network (RNN) with a many-to-many activation regime (Fig. 1). This configuration generates silence and vocalization activity class posteriors for each extracted feature vector in real-time. The trained system accurately detected 87.91% of zebra finch vocalizations, with an 11.54% false alarm rate, and only missed one vocalization (0.55%) in our test data, using a tuned confidence threshold (see methods for details). Upon detecting the correct vocalization (distance calls in this case), the system triggered the feeder. This closed-loop VAD system enabled us to effectively train the birds to use their distance calls to obtain food.

### Zebra finches learned to use their distance calls for food request

We trained 10 zebra finches, placing each bird in the training chamber twice daily (see methods for details). Initially, during the pre-training phase, the birds emitted very few distance calls, as these are typically used for communication with conspecifics. To encourage the birds to use distance calls during training, we employed distance call playbacks, which have been shown to prompt vocal responses ^30–33^. This helped the birds understand that emitting a distance call would open the feeder.

After training, 9 out of 10 birds successfully learned to use their distance call to request food (watch Supplementary Video S1). The rate of distance calls significantly increased only under food-restricted condition after training (Fig. 2a). A comparison of call rates in the restricted condition before and after training, showed a significant increase, supporting the effectiveness of training (n=9 birds, Friedman test, post-hoc comparison: p=0.0206). Additionally, after training, birds emitted more distance calls when they were food-restricted compared to when they were well-fed (n=9 birds, Friedman test, post-hoc comparison: p=0.0429). Full details on the distance call rates are available in Supplementary Table S1.

**Figure 2.**
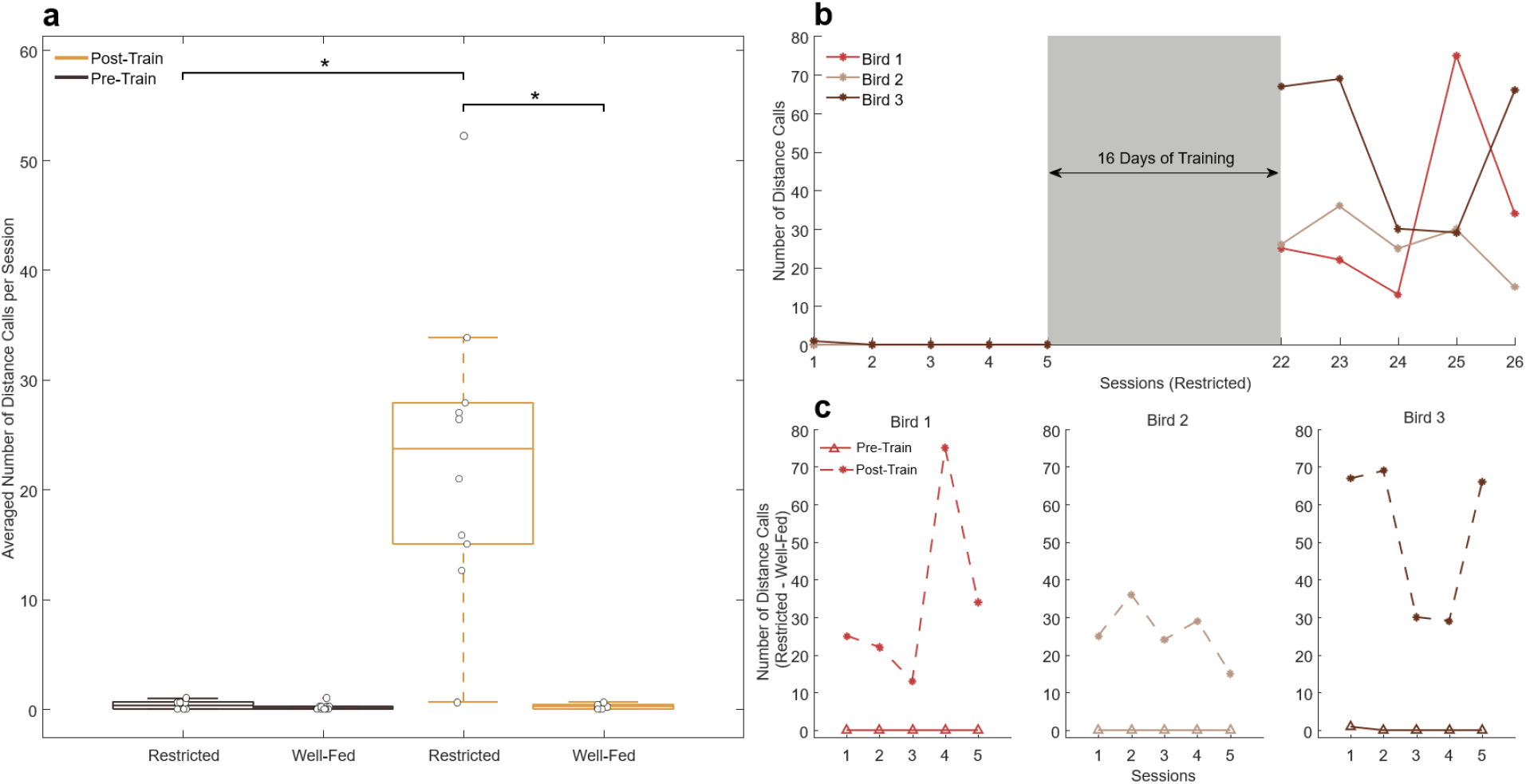
Birds learned to use their distance calls for food request. (**a**) Box plots showing the mean number of distance calls produced by the birds averaged over five sessions across four conditions; from left to right: Pre-train Restricted, Pre-train Well-fed, Post-train Restricted, and Post-train Well-fed. (**b**) The number of distance calls over time during restricted sessions for three example birds. Pre-training and post-training phases are shown, while the training phase is omitted due to the significantly higher distance call rates caused by playback of distance calls (see Methods). (**c**) Difference in the number of distance calls between food-restricted and well-fed conditions in pre-training and post-training for three example birds. Significant differences were assessed using the Friedman test, followed by post-hoc comparisons (*p < 0.05).

Before training, the call rate of the birds under food-restricted conditions was close to zero across five repetitions. After 16 days of training, the call rate for food requests increased and remained consistently high in five test repetitions (Fig. 2b). During the test phase, compared to the pre-training phase, the difference in distance call rates between food-restricted and well-fed conditions significantly increased (Fig. 2c), demonstrating that the zebra finches established a clear association between their distance calls and food requests. This data strongly suggests that this new novel paradigm was successful in learning the birds to use their distance calls to request food.

### Zebra finches can modulate the acoustic features of their calls based on their goal

During the final testing sessions, we noticed a distinct difference between the distance calls used in response to other zebra finches (audience-directed distance call, or aDC) and those used to request food (goal-directed distance call, or gDC). This observation raised the question of whether the birds could modulate the acoustic features of their calls depending on the context. To explore this, we conducted additional tests with two trained restricted birds (Bird 1 and Bird 2), in which a conspecific audience, visually separated from the subject, attempted to make contact with him.

We manually labeled the distance calls; if a distance call from the audience occurred within 3 seconds before the subject’s distance call, it was labeled as aDC; otherwise, it was labeled as gDC. Figure 3a shows the spectrogram of a 9-second example segment from the test session of Bird 1 (hear Supplementary Audio S1). The subject uttered a distance call without any preceding audience call (labeled as gDC) then the VAD correctly detected it and opened the feeder. Subsequently, the subject responded three times to two distance calls from the audience (labeled as aDCs). Figure 3b displays a zoomed-in view of the gDC, while fig. 3c shows the third aDC. Notably, the aDC has higher power and is more harmonic, with smoother high-frequency components.

**Figure 3.**
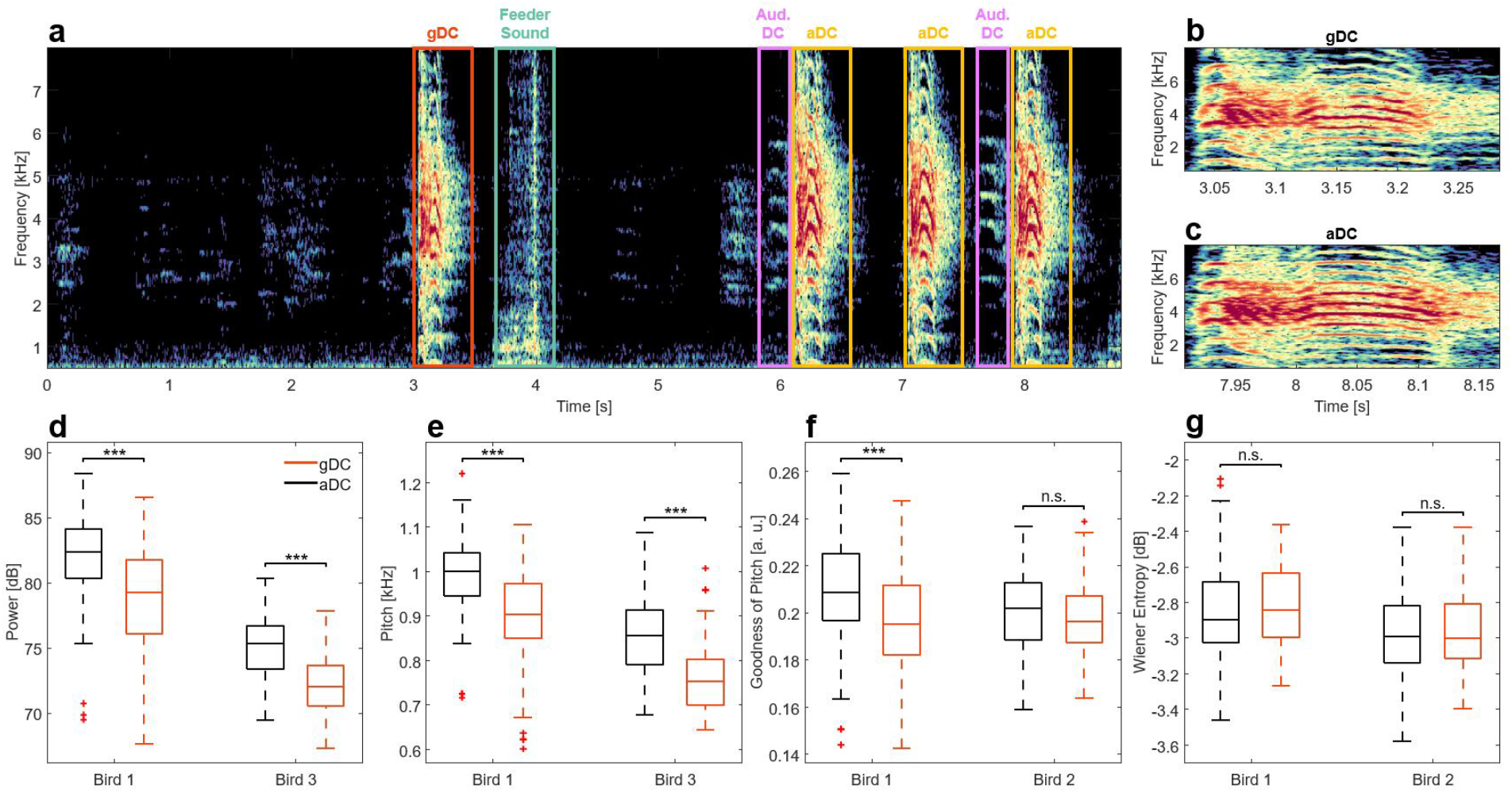
Acoustic flexibility of distance calls. (**a**) Spectrogram of a 9-second segment from one of the recordings of the acoustic flexibility analysis experiment, with different vocalizations labeled. The audio for this segment is available in Supplementary Audio S1. Note that after the first distance call the feeder opened and remained open for five seconds. This is why there is no feeder sound after next distance calls. (**b**) Zoomed-in spectrogram of the goal-directed distance call (gDC) shown in panel (a). (**c**) Same as panel (b), but for the third audience-directed distance call (aDC). Note the distinct differences in power and harmonic structure between the gDC and aDC. (**d-g**) Box plots comparing acoustic features of aDC and gDC in two birds: (**d**) Power, (**e**) Pitch, (**f**) Goodness of Pitch, and (**g**) Wiener Entropy. Significant differences were determined using the two-sample Kolmogorov–Smirnov test (**p < 0.01, ***p < 0.001). Abbreviations: gDC, goal-directed distance call; aDC, audience-directed distance call.

Our analysis exhibits significant differences in the acoustic features between aDCs and gDC in both birds. The distribution of vocalization powers differed significantly between the aDC and gDC (two-sample Kolmogorov–Smirnov test, Bird 1: n aDC=199, n gDC=131, p=9.3313e-11; Bird 2: n aDC=127, n gDC=72, p=9.9341e-13). Interestingly higher mean of aDC power suggests that the birds used louder vocalizations to reach distant audience. This is plausible given that the birds outside are farther away, and it has been shown that the auditory cortex of zebra finches can encode the distance of a propagated distance call ^34^. Additionally, the modulation of distance call amplitude aligns with prior findings showing that zebra finches can regulate the amplitude of their vocalizations depending on the social context ^35^.

Further analysis revealed that the distribution of vocalization pitch differed significantly between the aDC and gDC (two-sample Kolmogorov–Smirnov test, Bird 1: n aDC=199, n gDC=131, p=8.0166e-14; Bird 2: n aDC=127, n gDC=72, p=3.3964e-10). Previous elegant studies have shown that zebra finches can alter the pitch of their song syllables ^26,29,36^, as well as their distance calls depending on the internal state ^9^. We also found that the distribution of goodness of pitch differed significantly between the aDC and gDC, though this was only significant for one bird (two-sample Kolmogorov–Smirnov test, Bird 1: n aDC=199, n gDC=131, p=2.8021e-07; Bird 2: n aDC=127, n gDC=72, p=0.1295). Wiener entropy, another acoustic feature known to vary in corvids ^24^, did not show significant difference in our study (two-sample Kolmogorov–Smirnov test, Bird 1: n aDC=199, n gDC=131, p=0.3172; Bird 2: n aDC=127, n gDC=72, p=0.3971). Comparisons of other conventional acoustic features can be found in Supplementary Figure S1.

To further confirm acoustic difference between aDCs and gDCs, we used Linear Discriminant Analysis (LDA) to classify the vocalizations for each bird, using eight acoustic features (Power, Pitch, Goodness of Pitch, Wiener Entropy, Aperiodicity, Amplitude Modulation, Frequency Modulation, and Mean Frequency). The accuracies of the LDA classifiers were calculated using 5-fold cross-validation. The classification accuracy for bird 1 was 73.03% and for bird 2 was 82.91%. Figure 4a and 4b show the scatter plot of the correct and incorrect predictions of LDA classifier shown for the two most discriminative acoustic features which where power and mean frequency. It’s notable that the aDC and gDC had significant difference in these two features. Also, confusion matrices are presented in fig. 4a and 4b. Together, these results indicate that zebra finches modulate their distance calls with respect to the context, suggesting the acoustic flexibility of distance calls.

**Figure 4.**
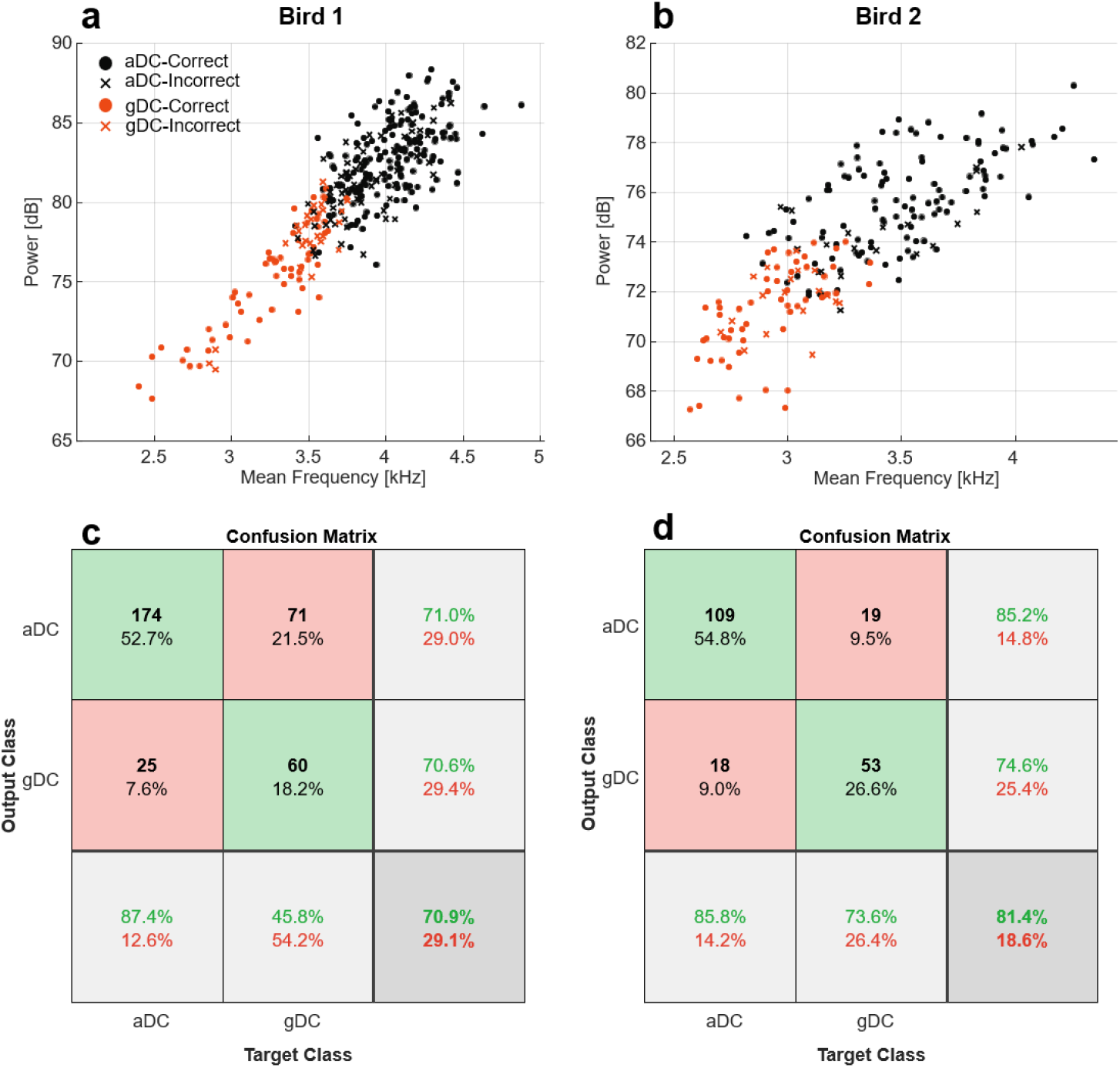
Classification of two distance call types. (**a**) Scatter plot of the correct and incorrect predictions of LDA classifier for Bird 1 shown for the two most discriminative acoustic features (power and mean frequency). (**b**) Same as panel (a) but for Bird 2. (**c**) Confusion matrix illustrating the classification accuracy of the linear discriminant analysis (LDA) model for Bird 1. (**d**) Same as panel (c) but for Bird 2. Abbreviations: gDC, goal-directed distance call; aDC, audience-directed distance call.

## Discussion

The transmission of information through vocal communication relies not only on the composition of vocal elements but also on the acoustic features of those elements. However, this form of information coding has not been extensively studied in animal models. To the best of our knowledge, we developed the first goal-directed vocalization paradigm in zebra finches, successfully training them to use distance calls to request food. More importantly, we indicated that zebra finches can modulate their vocalizations depending on context—distinguishing between goal-directed distance calls and those used for communication with conspecifics. Our results also align with the Convergent Pathways for Interaction (CPI) framework ^37^, suggesting that these modulations occur within the *State Processing* module, potentially facilitating communication of internal states and influencing motor programming.

The paradigm we developed offers exciting potential for future research, as it allows zebra finches to be conditioned under more complex situations. This is in some ways similar to a prior work conducted with Bengalese finches, where birds were shown to modulate their song under task-based conditions ^25^. For instance, in a preliminary test, we extended the paradigm to zebra finch songs, finding that the birds also learned to use songs for food request (Supplementary Video S2). Interestingly, they terminated songs uncompleted at syllable boundaries just after the feeder opened, which could provide valuable insight into the neural mechanisms underlying song initiation and truncation ^38^. Additionally, the paradigm could be employed to explore how zebra finches prioritize competing vocal behaviors, a key question in current research ^39^.

While we demonstrated the ability to modulate distance calls based on context, an important question remains: Can other birds perceive these subtle changes in vocalizations? Numerous studies have consistently shown that zebra finches are highly sensitive to fine acoustic variations ^40–43^. Zebra finches are capable of individual vocal recognition even if the bird has to distinguish males that all produce an imitation of the same song ^19^. They can even discriminate between different renditions of the same song, even down to individual syllables^44^. Moreover, they can use these fine variabilities to recognize different individuals in their group ^11–14,32,45,46^. Together all of these findings implicate that modulation of fine acoustic features can be perceived by zebra finches and can be used as a dimension in information communication.

Context-dependent vocal modulation is not a novel discovery. Many previous works demonstrated that the song in the contexts of courtship and territorial or aggressive encounters (“directed song”) is much more spectrotemporally ordered than songs produced in isolation or for daily training (“undirected song”) ^15,47,48^. Some previous works have also showed context-dependent modulations in calls; a famous study shows that chickadees alarm calls contain information about predator size ^49^. In zebra finches, vocal amplitude can be regulated based on social context ^35^ which we also observed in this study. Moreover, previous work have shown that some internal states like food availability can be reflected in acoustic features of the song ^50^. The ability to modulate the spectral features of distance calls has been previously reported ^9^. However, a key distinction in our findings is that while Perez et al. attributed this modulation to hormonal activity, our experiment indicates otherwise. Given that the aDC and gDC were vocalized in close temporal proximity, hormonal influence is unlikely. This suggests a capacity for moment-by-moment control over the acoustic features of vocalizations, implying that this modulation may be cognitively driven.

A further question arises: Is this ability to modulate acoustic features restricted to learned vocalizations? To our knowledge, the capacity to modulate spectral features has only been observed in learned vocalizations, such as songs and male distance calls. This raises the question of whether modulation is tied to the vocal learning pathway. Utilizing this paradigm with other calls of male zebra finches or even female zebra finches, who do not learn their vocalizations, could help answer whether they can modulate simpler features like amplitude too, or if more complex modulations are exclusive to learned vocal behaviors.

Previous studies were successful in categorizing different calls into semantic categories with fixed meaning ^51^ that are produced in an almost reflexive manner in a specific behavioral context ^52^. However, in this study, by developing a novel goal-directed vocalization paradigm, we trained zebra finches to use their distance calls for food request, raising the question of whether this training endowed the distance call with new meaning or simply established an association. Recent studies have shown the capacity to rearrange meaningless sounds in order to create new signals in some birds ^53–55^. In addition to compositional syntax, acoustic manipulations can also create new meaning as seen in the human language ^56,57^. To answer whether our paradigm have induced new meaning to distance call or not, further experiments should be conducted to see whether this usage of distance call for food request can be generalized to other contexts or even taught to other conspecifics. Importantly, distance call by itself lacks the semantics necessary for symbolic reference to external objects. However, our results can be interpreted in two ways. Firstly, this modulation of distance call may simply represent the bird’s internal state, without constituting a symbol. Alternatively, it can be viewed as a type of food call, potentially implying symbolic referencing ^58^. Additional research is required to clarify this.

It is also important to consider that the vocalizations in our experiment initiated without any cues. While hunger likely played a role, the initiation and modulation of distance calls suggest the possibility of volitional control. The idea that distance calls could be cognitively controlled is supported by studies showing that Bengalese finches can adjust the sequencing of vocal elements in response to learned contextual cues ^25,59^. Future research combining our task with existing paradigms used to explore volitional control over vocalization ^24^ could help address whether the observed modulations are indeed cognitively driven.

The neural circuitry involved in controlling vocal variability presents another exciting line of research. It is reasonable to hypothesize that circuits responsible for plasticity in song production are also involved in modulating distance calls. The anterior forebrain pathway, particularly the lateral magnocellular nucleus of the anterior nidopallium (LMAN), has been shown to regulate variability in song ^36^. Variability is transmitted to the vocal motor pathway through the robust nucleus of the arcopallium (RA) and HVC (proper name), both critical for song production and vocal learning ^60,61^. Interestingly, HVC and RA activity has been linked to food aversion, even in the absence of singing ^62^. It is also possible that different aspects of modulation, such as amplitude control versus spectral features, are governed by distinct circuits. This make sense since the pathways for learned and unlearned vocalizations are different ^63^. Because this type of context-dependent modulations need some form of cognitive control, it appears that caudal nidopallium (NC) which have been shown to be the equivalent of mammalian prefrontal cortex ^64^ is also involved. Recent work has shown that executive control of songbird vocalizations is done by NCL (the lateral region of NC) ^65^. At last, it seems that subcortical regions have also an important role in call initiation, especially the motor thalamic nucleus Uvaeformis (Uva) which its important role in driving vocal onsets have been shown recently ^38^. Altogether, the initiation and acoustic modulation of vocalizations may happen in a vast network including HVC, RA, LMAN, NC and Uva.

In summary, our study reveals compelling evidence that bird vocalizations are not just static and reflexive as previously thought. Instead, they can be dynamically modulated based on the context, showcasing a remarkable flexibility. This cognitive control over vocalization opens up new lines of research for studying animal communication. We hope that our goal-directed vocalization paradigm will offer valuable tools for further investigations. Also as proposed before ^66^, we believe that distance calls—given their learned properties and use in social contexts—offer unique opportunities to explore vocal communication and learning.

## Methods

### Subjects

Ten adult male zebra finches (*Taeniopygia guttata*), aged at least 120 days at the onset of the experiment, were sourced from the lab’s breeding colony. These males were housed in a mixed-sex aviary before and after the experiment. The birds were naïve to the experimental setup and training, and their social or genetic parentage, as well as genetic relatedness, was unknown. Throughout the experiment, the birds were maintained on a controlled feeding schedule and earned food during daily test sessions, with supplemental food provided as needed post-test. Water was provided *ad libitum* in both the aviary and test environments. The subjects were kept on a 12-hour light-dark cycle (lights on from 07:00 to 19:00).

### Apparatus

Training sessions were conducted in a custom-built wire cage (60 cm × 40 cm × 45 cm housed inside a sound-isolation chamber lined with sound-attenuating foam mats. The training chamber was illuminated by an overhead fluorescent lamp. The cage featured a single perch, and a camera with an integrated microphone was positioned on top to capture audio-visual recordings of the sessions. An online vocalization detection system was employed to monitor and classify the subjects’ vocalizations in real time. Upon detecting a distance call, the system activated an automatic feeder that delivered food pellets as a reward. Additionally, a speaker positioned above the cage was used to play auditory stimuli during training.

### Vocal activity detection system

Here we proposed a real-time vocalization detection and classification architecture for conditioning zebra finches to use their distance calls to obtain food. The proposed architecture consists of a one-stage detector/classifier recurrent neural network trained using the ^67^ dataset. The trained model is employed in the behavioral task of this study and can be further applied in other closed-loop experiments, potentially using other public or custom annotated datasets. The pipeline composition, data preparation, and model training are described below, with a quantitative evaluation of system performance provided in the results section.

The frame-based vocalization classifier is a left-to-right Long Short-Term Memory (LSTM) Recurrent Neural Network (RNN) that emits estimated vocalization class posteriors based on past and current acoustic features as follows:

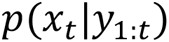

where (x_t_) is the vocalization class posterior, i.e., (p(x_t_ = [Sil, Distance, Distress, Tet, Thuk, Tuck, Whine] | y_1:t_) = [0.65, 0.06, 0.04, 0.13, 0.08, 0.04]); the current frame, (y_t_), is categorized as non-vocalization. The network is utilized and trained in a many-to-many operation mode, producing one posterior, (x_t_), for each new input (acoustic feature), (y_t_).

A total of 35 noise-augmented vocalization sequences were synthesized, each with a duration of approximately 720 seconds. For each sequence, selected target vocalizations from ^67^ were packed sequentially with randomized duration silences between two consecutive song/call vocalizations. Therefore, each sequence contains vocalization/silence parts. Both vocalization and silence parts may contain background noise, randomly chosen with randomized signal-to-noise ratios. Short-term discrete-time Fourier analysis was performed on all synthesized sequences with a window length of 23 milliseconds and a frame overlap of 11.5 milliseconds. Sixty linear filters were applied to each frame after applying a Hamming window, resulting in approximately 2 million feature vectors extracted as the training and validation dataset (augmented version of ^67^). Frame labels were selected from the silence and target vocalization set, relative to the frame’s correspondence to the silence or vocalization part.

A five-layer deep LSTM was trained in a many-to-many operation mode to assign each frame to a class label, implicitly performing two tasks simultaneously: vocalization detection and classification. The input dimension of the model is the same as the feature vectors, i.e., 60, and the model output contains 7 softmax neurons with cross-entropy loss to classify silence and other vocalization types (Distance, Distress, Tet, Thuk, Tuck, and Whine calls). Approximately 88% of the data (31 out of 35 sequences) were used for model training, with Adam used as the training algorithm. The trained model correctly classified 96.33% of the validation data.

For our task we partitioned calls either as “distance call” or “other calls”. Different from the ^67^ dataset we recorded a test data for validating the VAD system. The system accurately detected 87.91% of zebra finch vocalizations, with an 11.54% false alarm rate, and only missed one vocalization (0.55%) in our test data, using a tuned confidence threshold. This closed-loop VAD system was developed using MATLAB R2022b.

### Training procedure

Prior to the initiation of the experimental trials, the zebra finch subjects were acclimatized to the testing apparatus over a period of three days. This acclimatization phase was critical to mitigate any stress-induced variables that could potentially confound the behavioral outcomes. During this phase, the subjects were allowed to explore and become familiar with the confines of the device environment, thus reducing novelty-induced anxiety, which is known to affect performance in operant tasks. The birds did not receive any rewards during this phase. The experimental protocol consisted of three distinct phases: pre-training, training, and post-training.

1. **Pre-training**: Before initiating the conditioning protocols, a pivotal assessment was conducted to ascertain the correlation between the zebra finches’ vocalization patterns and their hunger levels. This phase was critical for establishing a control measure against which the efficacy of the operant conditioning could be evaluated. Additionally, this assessment ensured that the vocalizations were not a true reflection of the birds’ hunger state in innate behavior. Vocalization rates were measured within the apparatus under two distinct conditions: restricted and well-fed. This was performed five times for each condition. The order of restricted and well-fed experiments was randomized to prevent the response from becoming habitual due to repeating the same conditions. During the initial phase of food restriction, the zebra finches underwent a carefully regulated period of food deprivation lasting 5 hours. This systematic method of limiting food intake was precisely formulated to minimize any stress that could arise from abrupt food shortages. By gradually reducing food access, our objective was to ensure that the birds retained a sense of reliability in the availability of food, thus averting any negative impacts on their welfare or conduct.
2. **Training**: Subsequently, the subjects were placed in the apparatus twice a day in the morning and afternoon sessions. After each distance call, the feeder was opened. From the results obtained in the pre-training phase, the birds in the cage were usually silent; to enhance the learning process, playbacks of the distance call were employed as an auditory stimulus to provoke the subjects’ vocal responses, which in turn opened the feeder. Over time, the birds associated the distance call with obtaining food. During this period, the birds had no access to food outside the apparatus, ensuring they had enough motivation to learn the task. If the bird received an insufficient amount of food in the apparatus, additional food was provided a few hours after each training session to prevent weight loss.
3. **Post-training**: During this period, the birds did not hear any calls, and we investigated whether there was a correlation between the birds’ vocalizations and their level of hunger. Similar to the pre-training phase, the test was conducted under two conditions - restricted and well-fed. The order of these conditions was randomized, and each condition was tested in 5 sessions.

For the acoustic flexibility analysis experiment, in order to investigate our hypothesis (regarding the context-dependent acoustic flexibility of distance calls), we removed the sound attenuation chamber and introduced four conspecific audience members (both male and female) that were visually separated. The birds attempted to make contact using distance calls. This experiment was conducted on two of the trained birds when they were food-restricted.

### Data analysis

Labeling the recorded signal was done online by the VAD system, and all subsequent analyses were performed offline using MATLAB R2022b. The analysis aimed to determine whether the four experimental conditions—pre-train restricted, pre-train well-fed, post-train restricted, and post-train well-fed— produced significant differences in the number of distance calls in zebra finches. First, the data of the unlearned bird was removed. Given the small sample size (9 subjects) and the repeated measures design we utilized the Friedman test, a non-parametric alternative to the repeated measures ANOVA, to assess differences across the four conditions. If the Friedman test indicated a significant difference between conditions, post-hoc pairwise comparisons were conducted by Wilcoxon test.

For the acoustic flexibility analysis experiment, we labeled the distance calls manually. If a distance call from the audience occurred within three seconds before the subject’s distance call, it was labeled as aDC; otherwise, it was labeled as gDC. It is worth noting that the VAD system recognized both aDC and gDC as distance calls and opened the feeder, so the labeling was not on the base of feeder openings. Then we used SAP2011 ^68^ to extract acoustic features of distance calls. The extracted acoustic features were vectors with the same dimension as the distance call itself. The mean value of that vector was calculated so that every distance call had a single value for each of its acoustic features. To find significant differences between aDC and gDC, we used the two-sample Kolmogorov–Smirnov test because the assumptions of normality and homogeneity of variance were not met for all features.

Finally, for the classification, a linear discriminant analysis (LDA) classifier was trained on the data using MATLAB’s “fitcdiscr” function. We used 8 acoustic features (Power, Pitch, Goodness of Pitch, Wiener Entropy, Aperiodicity, Amplitude Modulation, Frequency Modulation, and Mean Frequency) as predictors. To assess the generalizability of the model, five-fold cross-validation was performed using MATLAB’s “crossval” function. The confusion matrices were calculated by running the trained models on the entire datasets.

### Ethical statement

All procedures were approved by the Research Ethics Committee of the School of Cognitive Sciences (SCS) of the Institute for Research in Fundamental Sciences (IPM, protocol number 1402/40/1/2841).

## Acknowledgements

We extend our deepest gratitude to Dr. Ali Ghazizadeh for his invaluable support in establishing the Birds Lab and for his insightful contributions throughout the experiments and manuscript preparation. We also wish to thank the IPM School of Cognitive Sciences for their essential support in creating the Birds Lab.

## Author Contributions

Z.S. conceptualized the study; All authors designed the methodology; M.K. created the VAD software; Z.S., P.M. and M.R. collected the data; A.B. and Z.S. analyzed the data; A.B., Z.S. and M.K. wrote the manuscript; All authors reviewed the manuscript; M.K. and M.R.R. supervised the study.

## Data Availability

The datasets for the current study are available from the corresponding authors upon reasonable request.

## Supplementary Information

**Supplementary Video S1.** This video showcases a 4-minute segment featuring one of the trained birds using its distance call to request food.

**Supplementary Table S1.** Detailed data showing the mean number of emitted distance calls averaged over five sessions across all four experimental conditions. Note the significantly higher values in the “Post-Train Restricted” column compared to the others. Bird #4, highlighted in red, represents the unlearned subject.

| Bird ID | Pre-Train<br>Restricted | Pre-Train<br>Well-Fed | Post-Train<br>Restricted | Post-Train<br>Well-Fed |
| --- | --- | --- | --- | --- |
| #1 | 0.6 | 0.2 | 15 | 0.2 |
| #2 | 0.6 | 0.2 | 12.6 | 0.4 |
| #3 | 1 | 1 | 21 | 0.2 |
| #4 | 0.6 | 0.2 | 0.6 | 0.2 |
| #5 | 0 | 0 | 33.8 | 0 |
| #6 | 0 | 0 | 26.4 | 0 |
| #7 | 0.6 | 0 | 15.8 | 0.6 |
| #8 | 0 | 0.2 | 52.2 | 0.4 |
| #9 | 0 | 0 | 27 | 0 |
| #10 | 0 | 0 | 27.9 | 0 |

**Supplementary Audio S1**. This audio file contains a 9-second sample segment from the acoustic flexibility analysis experiment. The spectrogram of this segment is shown in Fig. 3a. In this recording, the differences between the distance calls are distinctly audible, particularly in their power.

**Supplementary Figure S1.**
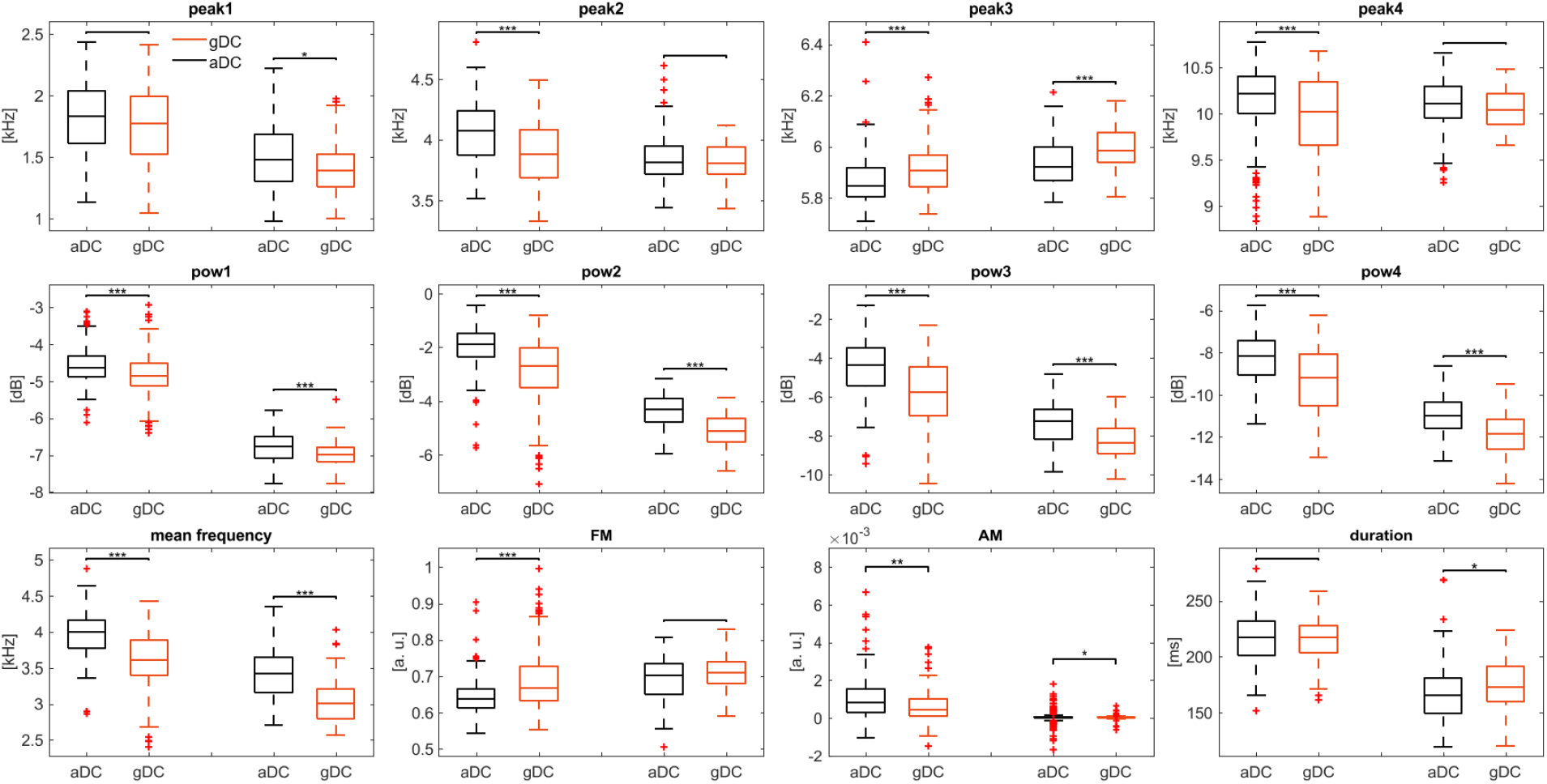
Box plots comparing the other acoustic features of aDC and gDC in two birds. Significant differences were determined using the two-sample Kolmogorov–Smirnov test (*p < 0.05, **p < 0.01, ***p < 0.001). Abbreviations: gDC, goal-directed distance call; aDC, audience-directed distance call.

**Supplementary Video S2.** This video showcases two birds from a pilot experiment that were trained to use their songs to request food. In this experiment, the closed-loop Vocal Activity Detection (VAD) system was calibrated to detect song vocalizations. Notably, both birds truncated their songs at the end of a syllable once the feeder was triggered and opened.

