## Supplementary figures and images for "Context-Dependent Modulations in Acoustic Features of Zebra Finch Distance Calls: Insights from a Novel Goal-Directed Vocalization Paradigm"

### Supplementary Figure S1

#### Slide 1
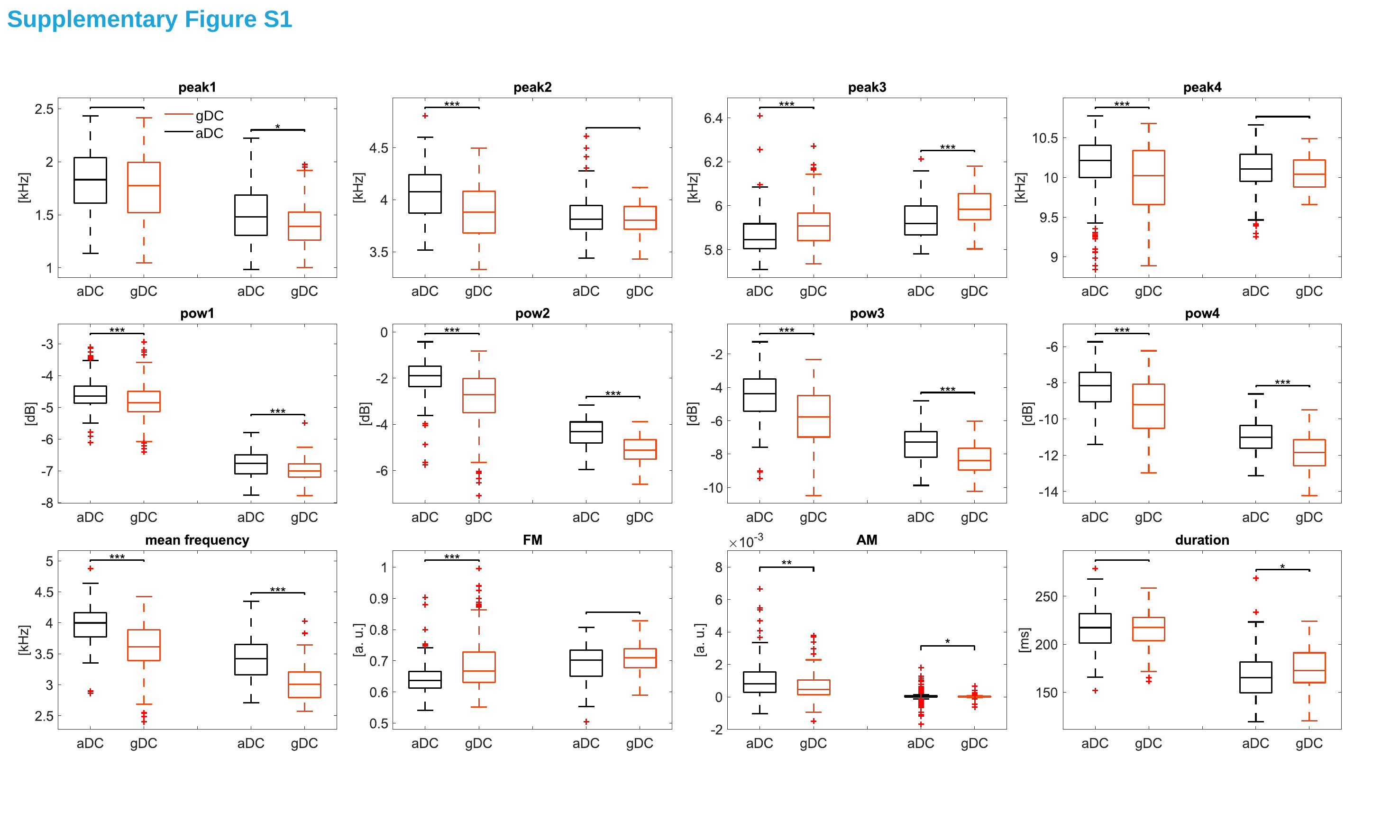

Supplementary Figure S1
gDC
aDC
